# The dynamic interactive network of long non-coding RNAs and chromatin accessibility facilitates erythroid differentiation

**DOI:** 10.1101/2021.03.02.433076

**Authors:** Yunxiao Ren, Junwei Zhu, Yuanyuan Han, Pin Li, Hongzhu Qu, Zhaojun Zhang, Xiangdong Fang

## Abstract

Erythroid differentiation is a dynamic process regulated by multiple factors, while the interaction between long non-coding RNAs and chromatin accessibility and its influence on erythroid differentiation remains unclear. To elucidate this interaction, we employed hematopoietic stem cells, multipotent progenitor cells, common myeloid progenitor cells, megakaryocyte-erythroid progenitor cells, and erythroblasts from human cord blood as an erythroid differentiation model to explore the coordinated regulatory functions of lncRNAs and chromatin accessibility in erythropoiesis by integrating RNA-Seq and ATAC-Seq data. We revealed that the integrated network of chromatin accessibility and LncRNAs exhibits stage-specific changes throughout the erythroid differentiation process, and that the changes at the EB stage of maturation are dramatic. We identified a subset of stage-specific lncRNAs and transcription factors (TFs) that coordinate with chromatin accessibility during erythroid differentiation, in which lncRNAs are key regulators of terminal erythroid differentiation via a lncRNA-TF-gene network. LncRNA *PCED1B-AS1* was revealed to regulate terminal erythroid differentiation by coordinating GATA1 dynamically binding to the chromatin during erythroid differentiation. *DANCR*, another lncRNA that is highly expressed at the MEP stage, was verified to promote erythroid differentiation by compromising megakaryocyte differentiation and coordinating with chromatin accessibility and TFs, such as RUNX1. Overall, our results identified the interactive network of lncRNAs and chromatin accessibility in erythropoiesis and provide novel insights into erythroid differentiation and abundant resources for further study.

**Key Points:**

- LncRNAs regulate erythroid differentiation through coordinating with chromatin accessibility.
- The integrative multi-omics analysis reveals stage-specific interaction network of LncRNAs and chromatin accessibility in erythropoiesis.

## Introduction

The human body produces approximately 2 x 10^11^ erythroid cells per day to meet the requirements for oxygen delivery and carbon dioxide exchange^1^. Erythroid cells are the main components of blood, healthy erythrocyte counts are essential for successful blood transfusions. In adults, the process of erythropoiesis is generated from hematopoietic stem cells (HSCs) in the bone marrow, which differentiate into multipotent progenitor cells (MPPs) and common myeloid progenitor cells (CMPs), followed by differentiation into megakaryocyte-erythroid progenitor cells (MEPs), erythroid progenitor cells and erythroblasts (EBs), which then de-nucleate to form reticulocytes that are released into the bloodstream where they eventually become erythrocytes^2,3^.

Erythroid differentiation is a dynamic, complex, and precisely regulated process. Any abnormal regulation may damage the formation of erythroid cells, which will cause erythropoiesis-related diseases. Previous studies have made advancements in elucidating the microenvironment and interaction of cytokines, transcription factors (TFs), microRNAs, and signaling pathways^1–6^, but the regulatory mechanisms of the interactivity of lncRNAs and chromatin accessibility lack comprehensive exploration.

The chromatin structure is essential for gene expression. Chromatin accessibility is enabled binding by chromatin regulators like TFs and cis-elements, which are involved in transcriptional regulation in eukaryotes^7–11^. Through the transcriptome and chromatin accessibility analysis, some studies discovered regulatory heterogeneity during early human hematopoiesis, which indicates a unique regulatory evolution in hematopoietic diseases^10–13^. However, few studies have focused on the interaction between long non-coding elements and chromatin accessibility during erythroid differentiation.

LncRNAs have a length greater than 200 nucleotides and are non-coding RNAs with a wide range of mechanisms that combine with DNA, RNA, and protein to regulate gene expression and modify chromatin formation^14–18^. Previous studies found that lncRNAs are involved in the regulation of hematopoiesis and erythroid differentiation^14–17^, and lncRNA expression in erythroid differentiation presents strong stage specificity and poor conservation in species^19–23^. However, the mechanism by which these lncRNAs regulate erythroid differentiation has not been well clarified.

To better understand the regulatory functions of chromatin accessibility and lncRNAs during erythroid differentiation, we performed an integrative analysis of the chromatin accessibility and transcriptome data. We established a comprehensive landscape of chromatin accessibility and transcriptomes, especially the dynamics of lncRNAs during erythroid differentiation. We observed that dynamic changes in lncRNAs, chromatin accessibility, and gene expression are stage-specific throughout the erythroid differentiation process. We further identified a lncRNA-TF-gene regulatory network that regulates terminal erythroid differentiation. Two lncRNAs, including *DANCR* and *PCED1B-AS1*, were functionally characterized as regulators of erythropoiesis in coordination with the TFs and chromatin accessibility. Our findings provide new insights into the dynamic interactive network of lncRNA and chromatin in human erythropoiesis, which may lead to the discovery of biomarkers for preventing or treating erythropoietic or hematopoietic diseases.

## Methods

### Cell models and data availability

We employed HSCs, MPPs, CMPs, MEPs, and EBs from human cord blood as a cell model of erythroid differentiation. The raw RNA-Seq data used in this study were obtained from the BLUEPRINT^24^ database (8th data release, 2016), which include EGAD00001000907, EGAD00001000911, EGAD00001000915, EGAD00001000919, EGAD0000100939, EGAD00001001140, EGAD00001001156, EGAD00001001169, EGAD00001001177, EGAD00001001186, EGAD00001001492, EGAD00001001515, EGAD00001001538, EGAD00001001550, EGAD00001001561, EGAD00001002316, EGAD00001002358, EGAD00001002363, EGAD00001002433, and EGAD00001002478. The ChIP-seq data were downloaded from the BLUEPRINT Data Coordination Center portal. The ATAC-seq data were derived from the GSE74912 dataset. RNA-Seq datasets for DANCR-overexpressed K562 cells are available in the GSA (Genome Sequence Archive), with accession number CRA003708 (https://bigd.big.ac.cn/gsa/s/YzxGU094).

### RNA-Seq data and ATAC-seq data preprocessing

FastQC was used to check the quality of RNA-Seq and ATAC-seq raw data. Trimmomatic and TrimGalore were used to remove the adapters and low-quality reads. RNA-seq reads and ATAC-seq reads were then aligned to the human genome (GRCh37/hg19) using STAR^25^ and bowtie2, respectively. Then, RSEM^26^ was used to quantify the transcripts on RNA-seq data, from which a gene expression matrix and transcriptional expression matrix were constructed.

Polymerase chain reaction duplication on ATAC-seq data was removed with Picard tools, and mitochondrial reads were removed with SAMtools. Peaks finally obtained using MACS2^27^.

The phantompeakqualtools package was used to calculate the chain cross-correlation^28^. Chain cross-correlation is an effective way to assess the quality of ATAC-seq or ChIP-seq that does not depend on the peaks obtained but on prior experiments^28^. Bioconductor package ChIPQC was also used to further evaluate the signal distribution of the ATAC-seq^29^.

### Functional enrichment analysis of differentially expressed genes

The raw counts matrix was filtered with rowSums equal to 0, and DESeq2 was used for differential analysis. *Padj* <0.05 and |log2FoldChange|>1 were used as the threshold (P adjust value, *Padj*). Then, the cluterProfiler R package^30^ was used to analyze and visualize functional profiles of gene and gene clusters from Gene Ontology and Kyoto Encyclopedia of Genes and Genomes databases.

### Weighted gene co-expression network analysis

Weighted gene co-expression network analysis (WGCNA) is performed to analyze gene expression patterns of multiple samples, by which genes with similar expression patterns can be clustered into one module, so that the correlation between the module and specific traits can be calculated, which facilitates the identification of key regulation factors and elucidates the mechanisms of biology development, tumorigenesis, and other diseases^31^. We constructed 14 gene co-expression modules of each lineage during erythroid differentiation using weighted gene co-expression network analysis (WGCNA R software package).

The connectivity of an intra-module network refers to the sum of the correlations among genes in that module. Genes with high connectivity within a module are considered to be hub genes. Hub genes are usually regulatory factors, and thus are located upstream in the regulatory network, whereas genes with low connectivity are usually present downstream in the regulatory network^31^. We selected the modules that were significantly correlated with each cell type and calculated the intra-connectivity among the modules. Then, the hub genes in each module were screened according to the threshold of the module with a connectivity greater than 0.8.

### Differential peaks analysis

To analyze the changes in chromatin accessibility during erythroid differentiation, we chose the DiffBind R package as the tool for differential peak analysis according to the requirements of the input files and statistical models^32^. DiffBind supports peak dataset processing, such as peaks that intersect and merge, and identifies statistically significant binding sites based on the binding affinity. DESeq2 was used here for differential peaks analysis based on DiffBind.

### Peak annotation and functional enrichment analysis

We used the ChIPseeker R package to perform peak annotation^33^. We first defined the range of a promoter as 1 kb upstream or downstream of the transcription start site. annotatePeak function was used to annotate peaks. We obtained the genomic distribution feature of the peaks and lncRNAs, (i.e., promoter, intron, or exon). We then annotated the adjacent genes of peaks and extracted all the genes in the promoter and distal regions as well as the annotated genes, then conducted functional enrichment analysis using the clusterProfiler R package^30^.

### TFs Motif analysis

Motif enrichment analysis of each stage and the differential peaks was performed using HOMER software. The command module “findmotifsgenome.pl” with the parameter “-len 8, 10, 12” was used to identify motif sequences with lengths of 8, 10, and 12 bp. The motif files that were obtained were then filtered based on a *p-value* < 1e-10 threshold. The corresponding transcription factor expression of the motif was also filtered according to the threshold of FPKM > 5. We identified 64 TF motifs enriched in each cell phase during erythroid differentiation and 43 TFs motif enriched in differential peaks. We then selected some TFs with more stage-specific and plotted them with ggplot2 in R.

### Identification of mRNA and lncRNAs

We extracted mRNAs and lncRNAs based on genome annotation files from ensembl. The parameters used for mRNA extraction are type = “gene”, gene_biotype = “protein_coding”. The parameters used for lncRNA extraction were type = “transcript”, transcript_biotype = “lincRNA”, “Sense_intronic”, “bidirectional_promoter_lncRNA”, “sense_overlapping”, “antisense_RNA” and “3prime_overlapping_ncrna”.

Clustering analyses of mRNAs and lncRNAs were performed using hclust. We then screened for significant lncRNAs that may regulate erythroid differentiation by integrating the specific lncRNAs that were expressed in each differentiation stage with the differentially expressed lncRNAs in adjacent stages and the lncRNAs intersecting in the accessible chromatin region. We constructed a diagram of the intersection of the three datasets using UpSet plot with TBtools.

### Establishment of lncRNA-TF-gene regulatory network

We combined the ChIP-seq/DNase I-seq/ATAC-seq data of EBs from the Cistrome Data Browser toolkit (http://dbtoolkit.cistrome.org) with the previously screened lncRNAs to predict regulation factors within 1 kb of the lncRNAs and hub genes^33,34^. Regulatory factors that corresponded to EBs with regulatory potential greater than 0.5 were retained. We then construct a potential lncRNA-TF-gene regulatory network by integrating the TFs, hub genes and lncRNAs identified before.

### Identification of cis-regulation elements

Promotor, enhancer, and insulator genes were annotated with the ChIP-seq data of H3K4me3, H3K27ac, and H3K9me3, respectively. The genome profile was visualized using Gviz R package^35^.

### Cell Culture

We isolated CD34^+^ cells from human cord blood using EasySep^™^ Human CD34 Positive Selection Kit II and tested for purity over 95% using flow cytometry. For erythroid differentiation, CD34^+^ cells were cultured at 10^5^ cells/ml for seven days in SFEM II medium, supplemented with 1% penicillin/streptomycin, 10 ng/ml stem cell factor, 10 ng/ml IL-3, and 3 IU/ml erythropoietin (EPO). Then, the cells were cultured at 4 x 10^5^ cells/ml for four more days in SFEM II supplemented with stem cell factor and EPO, and cultured at 7.5 x 10^5^ cells/ml for 3 more days in SFEM II supplemented with EPO. Human CB samples from full-term newborns were obtained from Beijing Obstetrics and Gynecology Hospital, Capital Medical University (Beijing, China) with parental consents. The subject has signed an informed consent form, and the research was approved by the ethical committee of Beijing Institute of Genomics (BIG), Chinese Academy of Sciences.

K562 cells were cultured in the RPMI-1640 medium containing 10% fetal calf serum (Gibco) and 1% penicillin/streptomycin and induced towards erythroid lineage with 50 µM hemin. A lentiviral overexpression system was used to overexpress DANCR in CD34^+^ and K562 cells^22^. Cells were cultured in an incubator at 37°C with 5% CO_2_.

### Hemoglobin concentration detection

The collected cell pellet was lysed with RIPA Lysis Buffer supplemented with 1 mM phenylmethyl sulfonyl fluoride, and the hemoglobin concentration was measured using a QuantiChrom^™^ Hemoglobin Assay Kit (BioAssay Systems) following the manufacturer’s instructions. The absorbance was measured with a microplate reader at OD400 nm.

### Flow cytometry

To conduct flow cytometric analysis, ~1 x 10^5^ cells were collected from the culture, centrifuged at 300 g for 5 minutes to remove the medium, and washed twice with 2 ml 1x Dulbecco’s phosphate-buffered saline (DPBS) (Gibco) solution containing 2% fetal bovine serum (Gibco) and 2 mM EDTA. After centrifugation at 300 g for 5 minutes to remove the supernatant, the cells were resuspended in 50 µl DPBS buffer and antibodies (anti-human CD71-PE and anti-human CD235a-APC) were added. Next, the cells were incubated at 4°C for 10 minutes in the dark. After incubation, the cells were washed twice with 1 ml of 1x DPBS buffer and resuspended in 200 µl DPBS buffer to prepare the cell suspension. A BD FACSAria II instrument was used for flow cytometric analysis, and the data analysis was performed with Flowjo software (Version 7.6, Three Star).

### Colony-forming unit assay

Colony-forming units (CFU) were generated in a cytokine-containing methylcellulose medium (MethoCult Media H4434, StemCell Technologies) by seeding 500 cells per well with 0.6 ml medium in a 12-well plate. After 12–14 days of culturing, multi-lineage colonies were counted under an inverted microscope.

## Results

### Dynamics of chromatin accessibility and transcriptome during erythroid differentiation

The number of peaks in chromatin accessibility reflect changes of the chromatin state. The average peaks of HSC, MPP, and CMP are similar, with more than 100,000 peaks each, while the number of peaks of MEP began to gradually decrease, then sharply decreased at EB with an average of only 33,457 peaks (Figure 1A). Chromatin accessibility condenses in terminal erythroid differentiation, and erythroid-specific TFs were active to participate in erythroid differentiation, such as GATA1 and KLF1^36–40^. The dramatically decreased peaks may be related to condensed chromatin and/or TFs binding. To verify whether the changes at the transcriptome level correspond with chromatin accessibility, we counted the number of genes and lncRNAs. The number of expressed genes in EBs were significantly lower than those at other differentiation stages (Figure 1B). The decreased number of expressed genes may be related to the process of terminal erythroid differentiation and development. However, interestingly, the number of lncRNAs in EBs increased significantly compared with those of adjacent stages, which is different from the changes in chromatin accessibility and expressed genes (Figure 1C). LncRNAs have been observed to act in a variety of ways, including binding to DNA and RNA, regulating TFs and chromatin structures^41–43^, and regulating terminal erythroid differentiation^19,41,44^. Moreover, the average number of lncRNAs in HSCs is also higher than those of the adjacent stages (Figure 1C). Previous studies also showed that lncRNAs hosted in HSCs are involved in the regulation of HSC stemness and differentiation^21,23^

**Figure 1.**
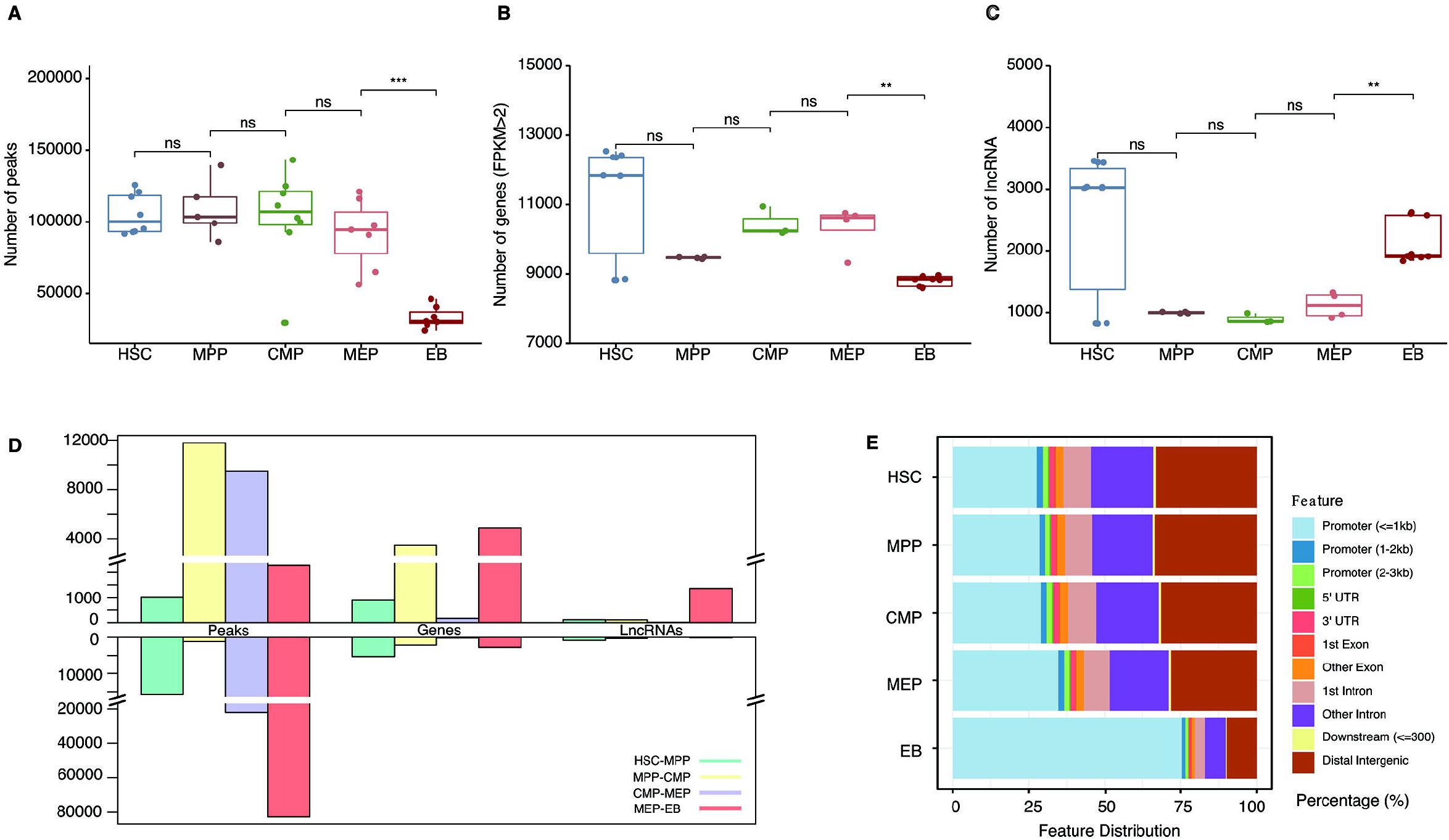
Dynamic changes in chromatin accessibility and the transcriptome during erythroid differentiation. (A) Peaks in ATAC-seq data. The boxplot illustrates the number of accessible chromatin regions at each stage during erythroid differentiation. (B) Number of genes at each stage from RNA-seq data with FPKM > 2. (C) Number of lncRNAs at each stage during erythroid differentiation. (D) Number of differentially accessible peaks, genes, and lncRNAs at adjacent stages during erythroid differentiation. The upper panel shows changes in the enhanced accessible chromatin, up-regulated genes, and lncRNAs of adjacent stages during erythroid differentiation, and the bottom panel displays the results of opposite changes. (E) Genomic distribution features of accessible chromatin. The bar plot shows the percentage of ATAC-seq peaks intersecting with the promoter, UTR, exon, intron, and distal intergenic. Statistical results were analyzed using a Kruskal Wallis test.

We further analyzed the significantly differentially accessible peaks, expressed genes, and lncRNAs of adjacent stages during erythroid differentiation. We found that the chromatin accessibility tended to decrease significantly during MEP differentiation into EBs (Figure 1D). There were more up-regulated lncRNAs than down-regulated lncRNAs, which exhibited a similar trend with the total number of lncRNAs between MEPs and EBs (Figure 1D). The genomic distribution of accessible chromatin showed that the proportion of accessible chromatin in promoters began to increase at the MEP stage and significantly increased at the EB stage by more than 75% (Figure 1E). This finding suggests that a large proportion of genes distributed in promoters may promote gene expression between MEPs and EBs, and the up-regulated genes involved in the terminal erythroid differentiation, including gradually removing the nucleus, endoplasmic reticulum, and mitochrondria, and then maturing into functional red blood cells that can transport oxygen^2^.

In addition, we observed a larger proportion of decreased accessible chromatin and up-regulated lncRNAs during HSC to MPP (Figure 1D), which is consistent with previous observations that epigenetic changes and lncRNAs have the potential to control the fate of stem and progenitor cells^23,45^. Another interesting finding is that there were more enhanced accessible chromatin and up-regulated genes at the CMP stage than at the MPP stage (Figure 1D). After this important divide in hematopoiesis, CMPs then differentiate into MEPs and granulocyte-macrophage progenitors (GMP)^46^. We found that there was a greater decrease in the accessible chromatin in MEPs than in CMPs, which indicated that the chromatin condensation starts during the MEP stage. However, there were few significant differentially expressed genes and lncRNAs between CMP and MEP (Figure 1D), which suggests that most are functionally conserved, while the significantly differentially expressed genes between them are supposed to promote erythroid differentiation from CMP to MEP. Taken together, dynamic changes in chromatin accessibility and lncRNAs were observed during erythroid differentiation, and the changes during the differentiation from MEP to EB stage were also dramatic.

### Characterization of functions associated with chromatin accessibility and the transcriptome during erythroid differentiation

To understand the biological functions of differential chromatin accessibility, differentially expressed lncRNAs and genes during erythroid differentiation, we performed a functional enrichment analysis. Compared to MEP, the enhanced accessible chromatins at the EB stage were enriched in the functions of erythroid differentiation (*p* = 3.18e-6), myeloid differentiation (*p* = 7.31e-7), actin cytoskeleton reorganization (*p* = 1.26e-5), and hemoglobin complex (*p* = 7.84e-5) (Figure 2A), which are essential processes during erythroid differentiation^47^. Consistently, the functions of up-regulated genes at the EB stage correlated closely with erythroid differentiation, including the functions of tetrapyrrole binding (*p* = 8.02e-6), heme binding (*p* = 2.95e-5), hemoglobin complex (*p* = 1.64e-13) and oxygen transport (*p* = 1.36e-10) (Figure 2B).

**Figure 2.**
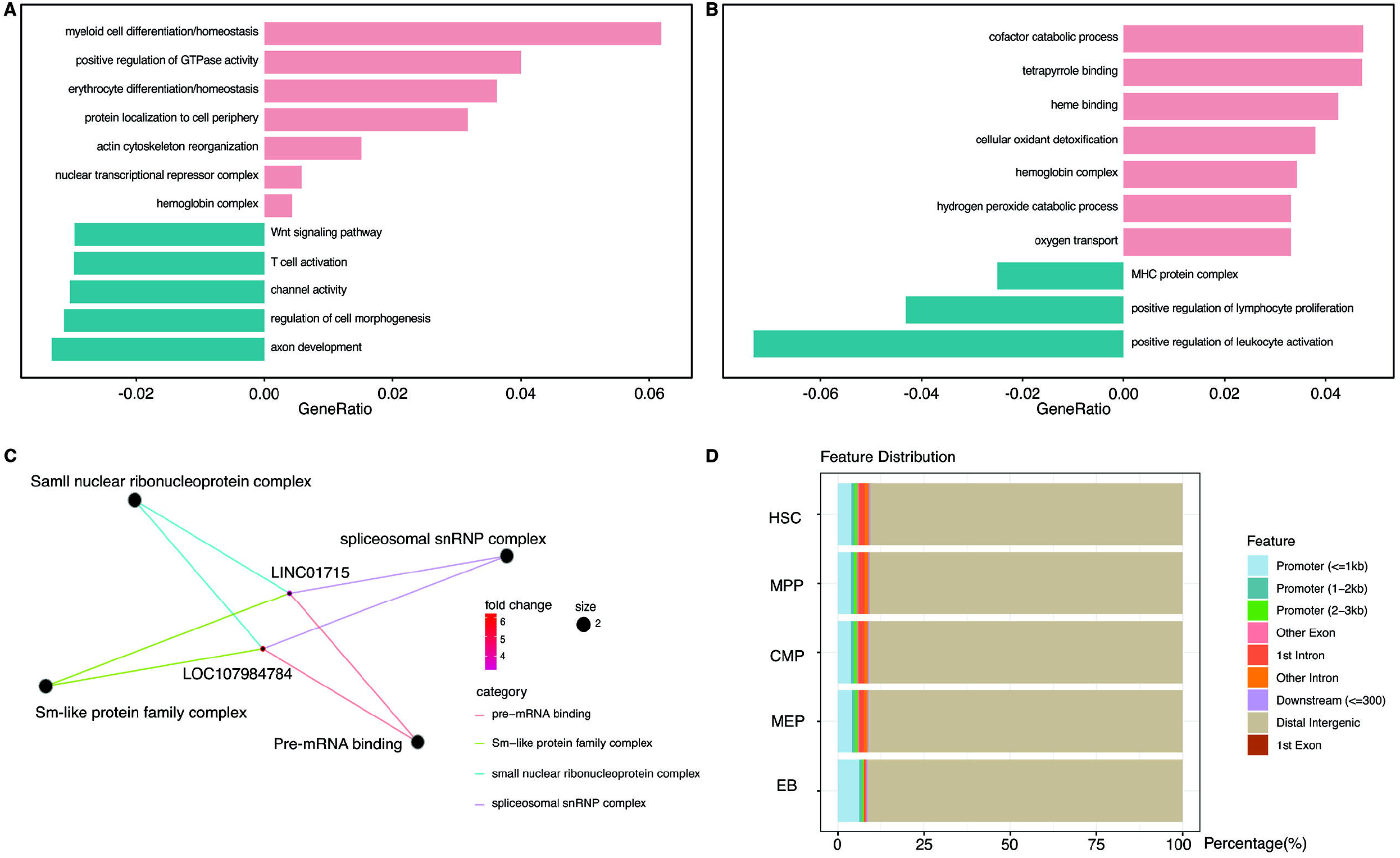
Function of differentially accessible chromatin and transcriptomes. (A-C) Functional enrichment of differential chromatin accessibility (A), differentially expressed genes (B), and lncRNAs (C) between the MEP and EB stages. (D) Distribution features of chromatin accessibility peaks related to lncRNAs.

Interestingly, the enrichment analysis of LncRNAs between MEP and EB showed that only two lncRNAs, *LOC107984784* and *LINC01715*, were annotated as relating to the functions of the spliceosomal small nuclear ribonucleoprotein complex (*p* = 0.69e-2) and pre-mRNA binding (*p* = 0.67e-3) (Figure 2C). To explore the relationship between lncRNAs and chromatin, we annotated the chromatin accessibility peaks on the lncRNA distribution features. We found that peaks related to lncRNA are mainly distributed on distal genomic regions (Figure 2D). The lncRNA located at the distal region may play a distal regulatory role by binding to enhancers or chromatin^48,49^.

In addition to the MEP and EB stage, we also analyzed the functions underlying the changes in chromatin accessibility and transcriptomes from HSC to MEP (supplemental Figure 1A). We noticed that enhanced chromatin accessibility of MEP was mainly related to myeloid and erythroid differentiation (*p* = 7.68e-8, *p* = 2.09e-07), and the up-regulated genes of MEP involve chromatin condensation (supplemental Figure 1 B), which indicates that the MEP period has begun in preparation for erythrocyte maturation.

### Stage-specific TFs and hub genes contribute to erythroid differentiation

Open chromatin regions can be bound with TFs and chromatin modifiers as an essential process in transcriptional regulation^50^. By integrating chromatin accessibility and the transcriptome profile in each stage, we performed motif enrichment analysis and characterized 64 TFs in each stage (Figure 3A, supplementary Table 1) and 43 TFs enriched in differential chromatin accessibility peaks (Figure 3B, supplementary Table2). Some TFs are well known to promote erythroid differentiation, like GATA1 and KLF1^51,52^, while function of the other TFs in erythroid differentiation is unclear.

**Figure 3.**
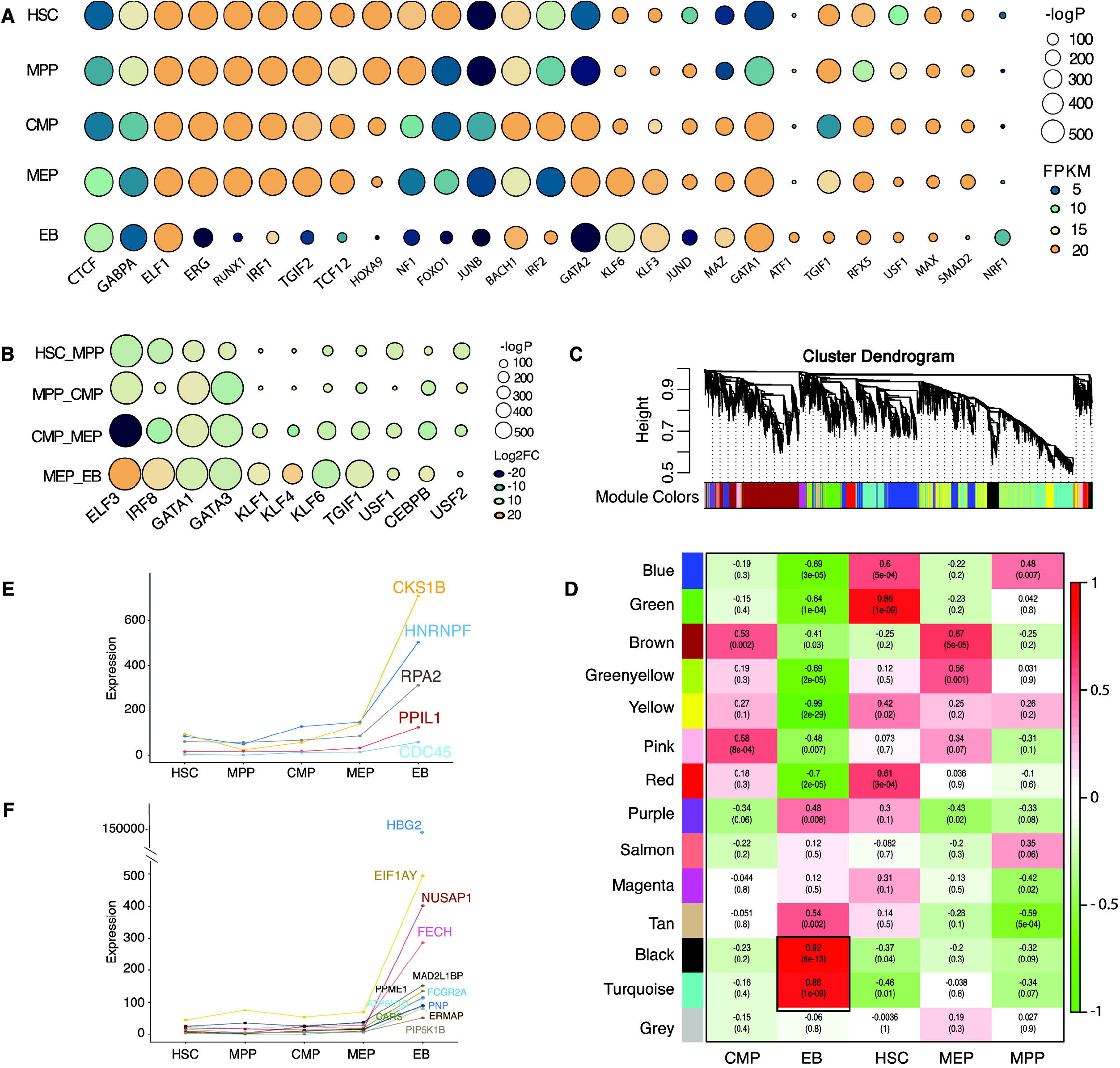
Transcriptional factors and hub gene identification through chromatin accessibility enrichment analysis and weighted gene co-expression network analysis. (A-B) TF motif enrichment of ATAC-seq peaks during erythroid differentiation. Panel A shows the TFs specifically enriched at each stage during erythroid differentiation; panel B shows TFs enriched at differential peaks. (Some of the TFs are shown in the figure according to their enrichment score). The size of the circle represents the -log *P-value*, which indicates the significance of TFs motif enrichment. The color of circle represents the expression (A, FPKM) and differential fold change (B, log2FC) of TFs. (C) Gene cluster dendrogram obtained through linkage hierarchical clustering. The colorful lines below the tree show the modules that were calculated by Dynamic Tree Cutting. (D) The relationships between module and cell type. Each row represents a module eigengene and each column corresponds to a trait. Each box contains the corresponding correlations and p-value. The colors in the figure are based on the correlations. (E-F) Expression of hub genes in EB, each shown by its significant module, the turquoise module (E) and the black module (F).

Hub genes refer to highly interconnected nodes in a module that plays significant roles in the regulatory network^31,53^. To further characterize the stage-specific hub genes which will help distinguish the regulatory networks underlying erythroid differentiation, we clustered genes with similar expression patterns into a module and identified 14 modules (Figure 3C). Each cell stage contains a significant co-expression module (Figure 3D). We selected the modules that were highly related to each stage and identified the hub genes specific to each module. A total of 5 and 12 hub genes were identified in two EB modules, turquoise module and the black module, respectively (Figure 3E-F). And 10, 14, 7, and 5 hub genes were also identified in the HSC, MPP, CMP, and MEP modules, respectively (supplementary Figure 2 A-E). The functions of some genes are well known, such as HBG2, which is characterized by the highest expression in EB stage. Two gamma chains combined with two alpha chains constitute fetal hemoglobin (HbF), which is normally replaced by adult hemoglobin (HbA) at birth^54,55^. However, some hub genes remain to be further explored. The identified stage-specific TFs and hub genes will be incorporated into the interactive networks during erythroid differentiation, which facilitates the regulatory mechanism underlying erythropoiesis.

### Chromatin-associated LncRNAs are involved in regulatory networks during erythroid differentiation

Hundreds of lncRNAs that promote erythroid differentiation and maturation are expressed specifically at each stage^19,56^. Our clustering results showed that lncRNA had stronger cell specificity compared with mRNAs (Figure 4A, B). We then extracted stage-specific lncRNAs during erythroid differentiation and displayed the top 10 lncRNAs with specific expression at each stage on a heatmap (Figure 4C). The differential lncRNA analysis showed a larger proportion of down-regulated lncRNAs from HSC and MPP to CMP (Figure 4D), while the proportion of up-regulated lncRNAs increased suddenly in EB (Figure 4D), which indicates that lncRNA may play an important role at the late stage of the erythroid differentiation.

**Figure 4.**
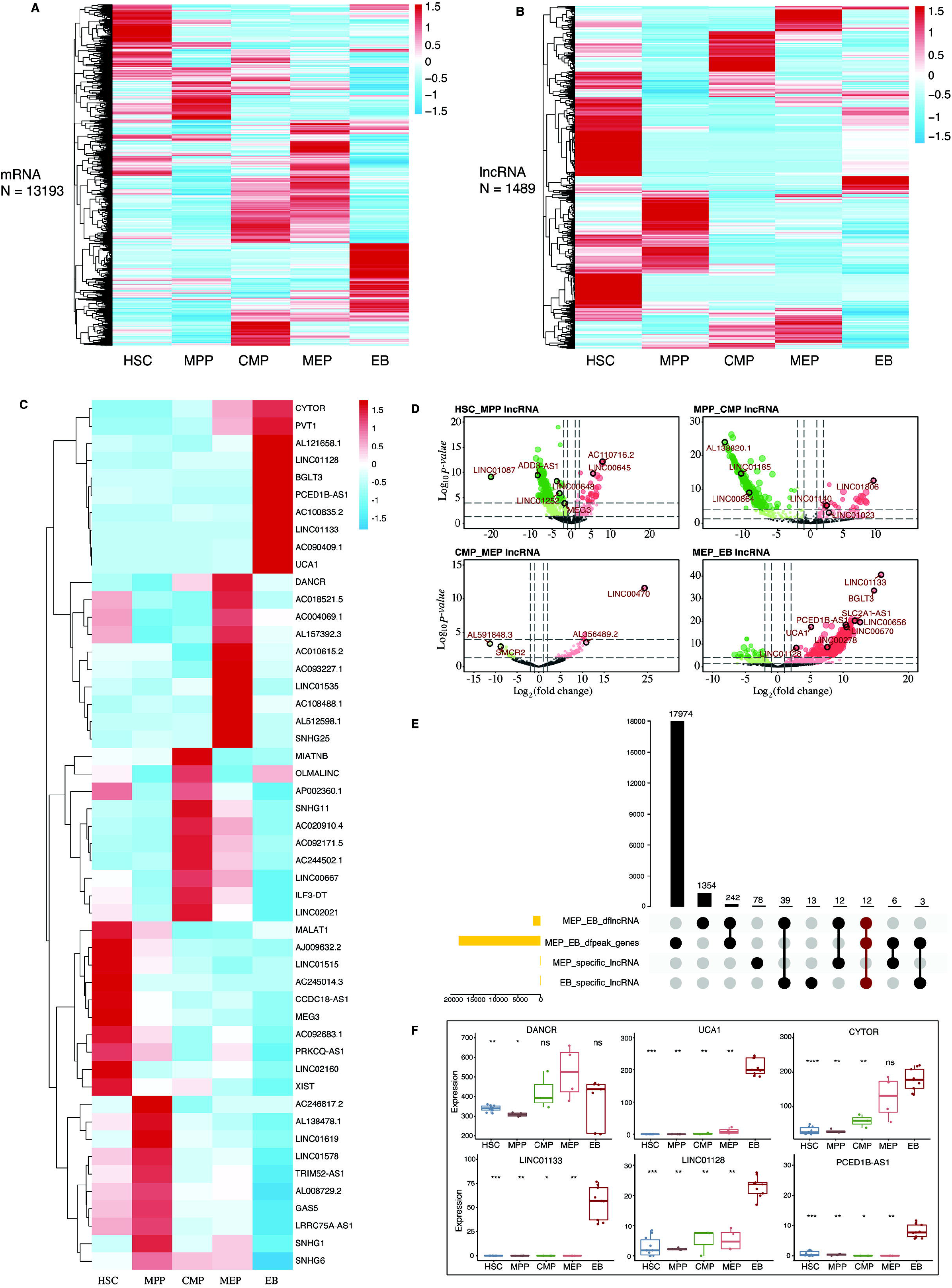
Identification of potential regulatory lncRNAs by integrating transcriptome and chromatin accessibility during erythroid differentiation. (A-B) Clustering results of mRNAs (A) and lncRNAs (B). (C) Top 10 stage-specific lncRNAs expressed during erythroid differentiation. (D) LncRNAs differentially expressed between adjacent stages during erythroid differentiation. (E) The number of intersecting differentially expressed lncRNAs and the accessible chromatin between the MEP and EB stage. The left bar plot with yellow color shows the number of differentially expressed lncRNAs and genes between MEP and EB, and the number of specific lncRNAs in EMP and EB. The bottom right dot plot connected by solid lines represent different intersecting points. The top right bar plot represents the number of intersections. (F) The expression of lncRNAs with potential regulatory functions at the MEP and EB stages. DANCR are highly expressed at MEP stage. *UCA1, CYTOR, LINC01133, LINC01128*, and *PCED1B-AS1* are highly expressed at EB stage.

Accessible chromatin can bind by cis-regulatory elements, such as enhancer^8,57^. We hypothesized that lncRNAs located in the distal region may interact with enhancer or chromatin to regulate erythroid differentiation. To test this hypothesis, we screened for lncRNAs that are differentially expressed, located in the differential accessible chromatin region, stage-specific, and highly expressed (Figure 4E, supplementary Figure 3 A-C). We identified 4, 3, 2, 1, and 5 lncRNAs at the stages of HSC (supplementary Figure 4 A-E), MPP (supplementary Figure 4 F, G), CMP (supplementary Figure 4 H, I), MEP (Figure 4F) and EB, respectively (Figure 4F), that could be associated with erythropoiesis, of which *DANCR* is specifically highly expressed in the MEP stage and was further functionally verified in this study.

### *DANCR* promotes erythroid differentiation by compromising megakaryocyte differentiation

We found that chromatin accessibility and the transcriptome changed dramatically at the MEP stage which can give rise to megakaryocytes and erythroid cells. We observed *DANCR* were specifically expressed higher at the MEP stage (Figure 4F). There are strong H3K4me3 and H3K27ac signals around the *DANCR* (Figure 5A), which indicates that this region has enhancer signal^58^. *DANCR* is a tumor promoter, however, little is known about its function in erythroid differentiation^59^. We observed that *DANCR* promotes gene expression of erythroid-specific globin genes (ε, γ- and β-globin) and *ALAS2* in DANCR-overexpressed K562 cells after 3 days induction towards erythroid lineage (supplemental Figure 5A, B). *DANCR* overexpression also increases the expression of hemoglobin in differentiated K562 and CD34^+^ cells (Figure 5C, supplemental Figure 5 C). The double-positive cells of CD235a^+^CD71^+^ in differentiated CD34^+^ cells and K562 cells with *DANCR* over-expressed were significantly higher than in the controls (Figure 5B, supplementary Figure 5D, E). These results illustrate that *DANCR* promotes erythroid differentiation. The transcriptome analysis of *DANCR*-overexpressed K562 cells showed that *DNACR* also inhibits megakaryocytes differentiation (Figure 5D). Colony-forming unit assay revealed that overexpressed *DANCR* significantly promotes the production of erythroid progenitors but inhibits megakaryocyte progenitor cells (*p* < 0.05), with myeloid progenitor cells unchanged, suggesting that *DANCR* promotes erythroid differentiation by compromising megakaryocyte differentiation (Figure 5E). To better understand the regulatory mechanism of *DANCR* in hematopoiesis, we used public ChIP-seq data^60^ to identify TFs that appear in the *DANCR* genomic region associated with chromatin accessibility (Figure 5F, supplemental Table 3), of which RUNX1 has a key role in regulating the balance between erythroid and megakaryocytic differentiation through modulating the balance between KLF1 and FLI1^61^. These results suggest that *DANCR* may coordinate TFs associated with chromatin accessibility in regulating MEPs differentiation.

**Figure 5.**
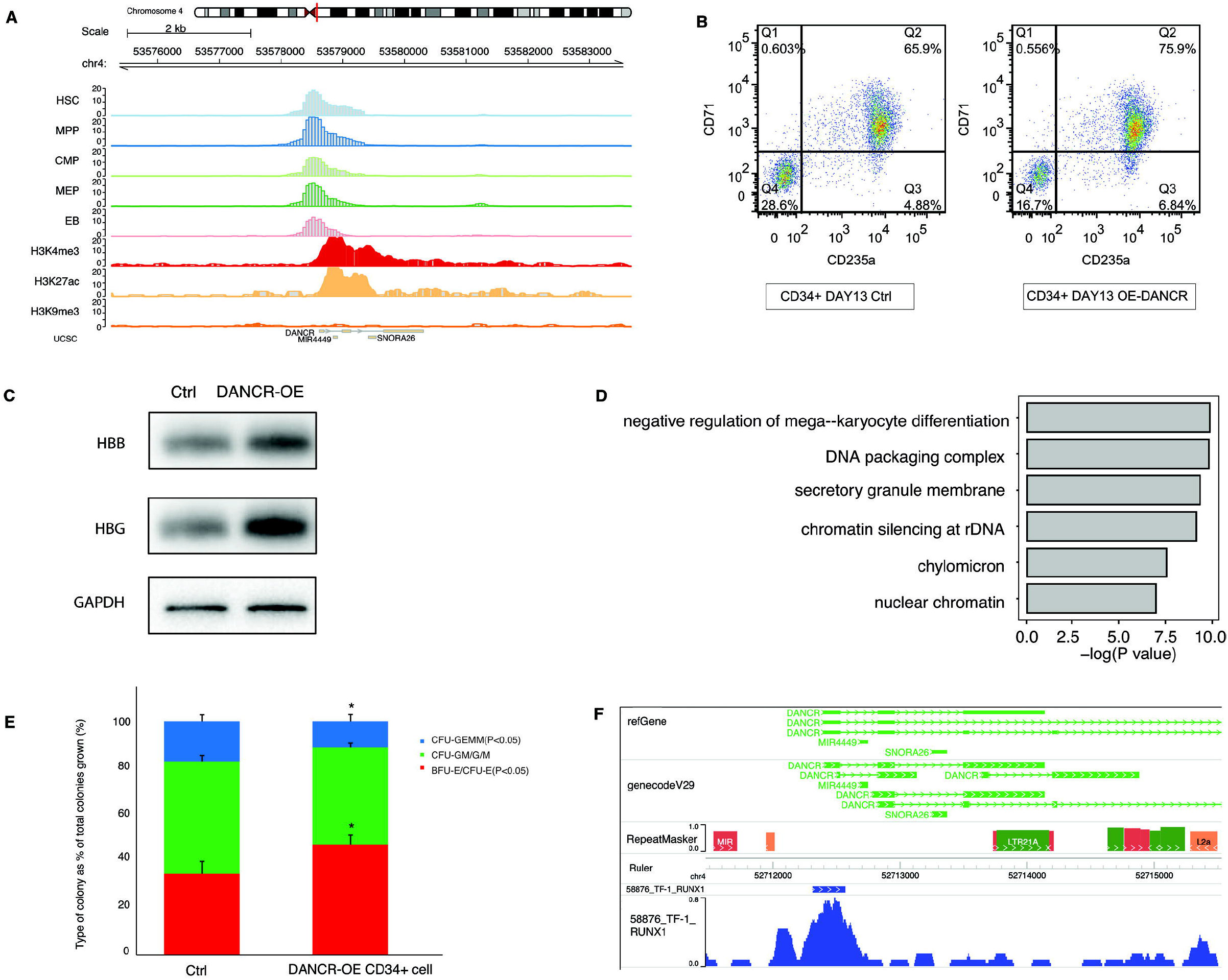
LncRNA *DANCR* is involved in erythropoiesis. (A) Chromatin accessibility and histone modification profile around *DACNR* locus. (B) The expression of CD235a and CD71 were detected by flow cytometry in the differentiated DANCR-OE CD34^+^ cells for 13 days. The percentage in the upper right of each figure represents the proportion of CD235a^+^-CD71^+^ double-positive cells. (C) Overexpression of DANCR in CD34^+^ cells promote the accumulation of β-hemoglobin (HBB) and γ-hemoglobin (HBG) proteins on Day 13. (D) Gene ontology enrichment of differential genes between DANCR-OE and control on day 0 in K562 cells. (E) Colony-forming capacity analysis of DANCR overexpression in CD34^+^ cells. The percentage of erythroid progenitor colonies (BFU-E/CFU-E, red) in DANCR-OE CD34^+^ cells was higher than that of the control, while the percentage of colonies (CFU-GEMM, blue) containing megakaryocyte progenitors in DANCR-OE CD34^+^ cells was lower than that of the control. (F) Profile of transcription factor RUNX1 around DANCR in TF-1 cells. Statistical results were analyzed by student’s t-test and Kruskal-Wallis test, *p < 0.05, **p < 0.01, ***p < 0.001. Ctrl: control group, OE: overexpression.

### Regulatory networks of lncRNAs, TFs, Genes involved in terminal erythroid differentiation

EBs are the important stage of erythroid differentiation involving the expulsion of the nucleus, which forms reticulocytes that mature into biconcave red blood cells^62^. We also found dramatic changes in chromatin accessibility and transcriptome occur at the EB stage (Figure1D): stage-specific TFs and genes are proposed to be involved in erythroid differentiation, which coordinates with changes in chromatin accessibility; lncRNAs expressed significantly higher at EB stage than at other stages. Thus, we hypothesize that there may be a regulatory network of lncRNAs, TFs, and genes that regulate terminal erythroid differentiation. We integrated public data including the ChIP-seq, DNase I-seq, and ATAC-seq data in Toolkit from Cistrome Data Browser to predict the potential regulatory networks at the EB stage (Figure 6A). The potential interactions are defined by cis and trans-regulation^34,63,64^.

**Figure 6.**
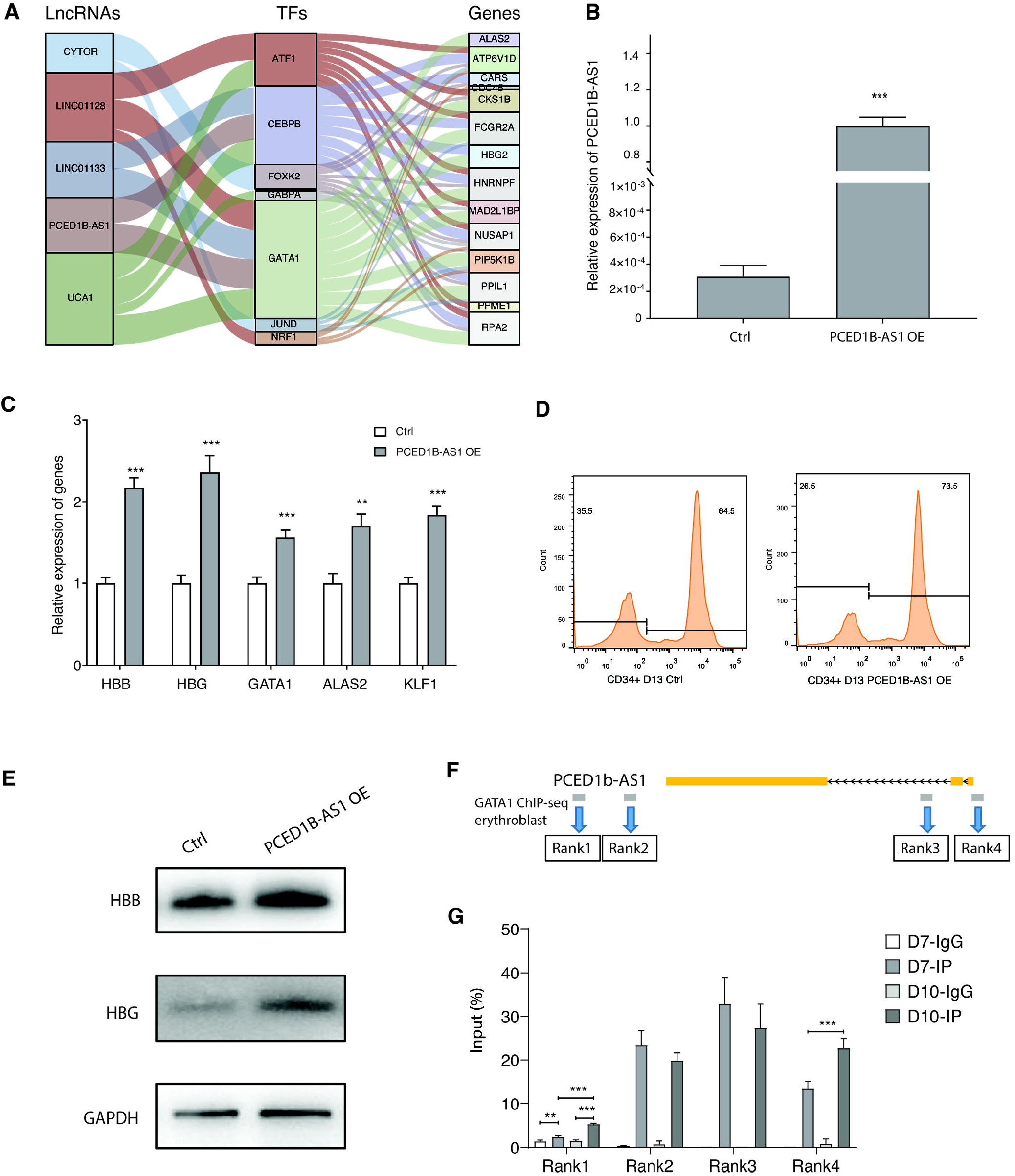
The regulatory networks of lncRNAs-TFs-Genes during terminal erythroid differentiation. (A) The lncRNAs-TF-gene regulatory network in EB. The left column in the figure shows the specific lncRNA identified in EB; the middle column shows the key TFs expressed in EB; and the right column shows the hub genes in EB. The line between the columns represents the possible regulatory relationship among factors. (B) The detection of *PCED1B-AS1* overexpression in differentiated CD34^+^ cells (Day 11) by qPCR assay. (C) Relative expression of erythroid related genes in differentiated *PCED1B-AS1-OE* CD34^+^ cells (Day 11) detected by qPCR assay. (D) The detection of CD235a^+^ cells by flow cytometry in the *PCED1B-AS1-OE* differentiated CD34^+^ cells (Day 13). The percentage on right in each figure represents the proportion of CD235a^+^ cells. (E) The expression of β-hemoglobin (HBB) and γ-hemoglobin (HBG) proteins detected by western blot in differentiated *PCED1B-AS1-OE* CD34^+^ cells (Day 13). (F) GATA1 binding sites around PCED1B-AS1 locus in EB stage from ENCODE database (ENCFF957CWW). The binding sites are named Rank1-4 from left to right. Rank3 locates in PCED1B-AS1 locus, the other 3 binding sites locate outside the PCED1B-AS1 locus. (G) GATA1 ChIP-qPCR analyses of IgG and IP on Day7 and Day10. The GATA1 binding signal was detected at the 4 regions (Rank1, Rank2, Rank3 and Rank4). Statistical results were analyzed by student’s t-test and Kruskal-Wallis test, *p < 0.05, **p < 0.01, ***p < 0.001. Ctrl: control group, OE: overexpression.

As shown in Figure 6A, LncRNA *UCA1* may coordinate with specific TFs, such as GATA1, ATF1, CEBPB, and GABPA, to regulate the expression of genes, including *ALAS2, HBG2*, and *ATP6V1D* (Figure 6A), which participate in terminal erythroid differentiation. Previously, one study reported that *UCA1* whose promoter is occupied by GATA1 functions as a scaffold lncRNA to maintain the stability of *ALAS2* mRNA for heme synthesis^65^, which is one crucial process for globin biosynthesis during erythroid differentiation. Our study found there is a significant increase on chromatin accessibility in the promoter region of *ALAS2* (supplemental Figure 6A). The *ALAS2* promoter region has no H3K9me3 signal, but has a weak H3K4me3 signal and a strong H3K27ac signal, suggesting that UCA1 may act as an enhancer lncRNA contributing to the binding of GATA1 and other regulatory factor complexes to *ALAS2* mRNA, which in turn regulates heme synthesis^65–69^.

*PCED1B-AS1* is another novel LncRNA that is specifically expressed at the EB stage. Our results showed that *PCED1B-AS1* overexpression promotes the expression of erythroid-related genes, including *HBB, HBG, GATA1, ALAS2* and *KLF1*, in erythroid cells differentiated from CD34^+^ cells (Figure 6B, C). Flow cytometry analysis showed that the proportion of CD235a positive cells was also increased (Figure 6D). *PCED1B-AS1* overexpression also increased the protein expression of HBB and HBG (Figure 6E). These results indicated that *PCED1B-AS1* promotes erythroid differentiation. Based on the regulatory network, we hypothesize that *PCED1B-AS1* regulates terminal erythroid differentiation cooperating with GATA1. Previously, we revealed the binding of GATA1 in the genomic regions of *PCED1B-AS1* in K562 cells^22^. In this study, using public ChIP-Seq dataset, we further screened four binding sites of GATA1 in genomic regions of *PCED1B-AS1* in erythroblasts, and revealed the binding ability of GTAT1 in two regions (Rank1, Rank4) gradually increased during erythroid differentiation of CD34^+^ cells (Figure 6F, G), demonstrating that *PCED1B-AS1* regulates terminal erythroid differentiation coordinating GATA1 dynamically binding to the chromatin during terminal erythroid differentiation. The regulatory networks of the other LncRNAs, including *CYTOR, LINC01128*, and *LINC01133*, with their TFs and genes were identified for the first time, which need to be further verified for their role in terminal erythroid differentiation. Taken together, lncRNAs located in the open chromatin region can serve as an enhancer lncRNA to recruit TFs or directly coordinate TFs and participate in erythroid differentiation.

## Discussion

In this study, our findings provide a comprehensive landscape of chromatin accessibility, lncRNAs, and hub genes, as well as trans-factors at each stage during erythroid differentiation, and identifies the interactive network of lncRNAs and chromatin accessibility in erythropoiesis, which provides novel insights into erythroid differentiation and abundant resources for further study. We modelled the interactive network of lncRNAs, TFs, genes in terminal erythroid differentiation and illustrated that several novel LncRNAs are probably involved in terminal erythroid differentiation cooperating with TFs, which provide new regulatory insights for erythropoiesis.

HSC and MPP are progenitor cells of erythroid differentiation with differentiation ability^70^. Chromatin accessibility tends to decrease from HSC differentiated into MPP as the expressed genes decrease. Some studies have reported that lncRNAs regulate differentiation and proliferation of HSC, such as *H19* and *MEG3*^21,71^. Interestingly, our results also indicate that down-regulated genes in HSC are associated with non-coding RNA processing. We identified lncRNA *CCDC18-AS1* was highly expressed at HSC. Studies showed that *CCDC18-AS1* involved in the cell cycle is similar to *MALAT1, NEAT1*, and *H19*^72^. Few studies have been conducted on *LINC01252* and *LINC00648*, which are the other two lncRNAs that are highly expressed specific to HSC. The proportion of enhanced accessible chromatin and up-regulated genes is larger during MPP differentiated into CMP (Figure 1A, B, D), which suggests that transcriptional regulatory activity remains active during MPP differentiated into CMP, and transcriptional regulators can bind to regulate this process.^45,73^

The MEP stage involves a continuous transition from CMP, in which the cells are bipotent and can further generate two completely different functional cells: erythrocytes and platelets. Our results show that the chromatin accessibility changes significantly, with numerous decreased accessible chromatin, which indicates that the MEPs prepare to or already possess some characteristics of mature erythroid cells. The functional annotation of changed chromatin accessibility and transcriptome illustrates their association with the functions of myeloid and erythroid differentiation. The differentiation fate of MEPs depends not only on TFs but also their target genes. Our findings expand the regulation mechanism of MEP differentiation and verify that *DANCR* can promote erythroid differentiation by compromising megakaryocyte differentiation by coordinating TFs. However, the specific target genes that are involved in this process require further verification.

In addition, we identified many novel hub genes and TFs at each stage (Figure 3) that may play important roles during erythroid differentiation, which could facilitate the understanding of the molecular networks underlying erythropoiesis. Importantly, we discovered a cluster of lncRNAs that are known or novel that play significant roles and interact with TFs and genes during erythroid differentiation. It has been reported that *H19* and *MEG3* regulate the differentiation and proliferation of HSC^21,71,74^. We also identified that *MEG3* is specifically expressed at the HSC stage.

EB is the transition stage between MEP and enucleated cells. Chromatin accessibility, lncRNA, and gene expression undergo tremendous changes during the differentiation of MEP into EB. Based on the previous study^22^, we further demonstrated, *PCED1B-AS1* regulates erythroid differentiation associated with GATA1 and chromatin remodeling in primary cells. *UCA1* may be involved in erythroid differentiation by recruiting TFs to target genes and chromatin state changes^65^. Our results identified the interactive network of lncRNAs and chromatin accessibility in erythropoiesis, and the functions of *CYTOR, LINC01128*, and *LINC01133* remained to be further elucidated in terminal erythroid differentiation.

Overall, our study characterized the interactive network of lncRNAs and chromatin accessibility during erythroid differentiation by multi-omics integrated analysis. We provide new perspectives and a rich resource for exploring the regulatory mechanism underlying erythroid differentiation, and offer the potential markers for preventing or treating various erythropoiesis-related diseases as well.

## Acknowledgments

This research was supported by the Strategic Priority Research Program of the Chinese Academy of Sciences (XDA16010602), National Natural Science Foundation of China (82070114, 81870097, 81670109, 81700097, 81700116), and National Key R&D Program of China (2017YFC0907400). This study makes use of data generated by the Blueprint Consortium. A full list of the investigators who contributed to the generation of the data is available from www.blueprint-epigenome.eu. Funding for the project was provided by the European Union’s Seventh Framework Program (FP7/2007-2013) under grant agreement no 282510 – BLUEPRINT.

## Authorship Contributions

XF and ZZ conceived and supervised the study; YR analyzed the data; ZZ and HQ designed the experiments; JZ, YH, and PL performed the experiments; YR drafted the manuscript; XF and ZZ revised the manuscript. All authors read and approved the final manuscript.

## Conflict of Interest Disclosures

The authors declare no competing financial interests.

## Additional Figure files

**Supplemental Figure 1.** Enriched function of differentially accessible chromatin (A) and expressed genes (B) during HSC differentiated into CMP.

**Supplemental Figure 2.** Expression of hub genes in significant modules of each stage during HSC differentiated into MEP.

**Supplemental Figure 3.** Integrative screening of lncRNAs. The upset plot shows the number of intersecting lncRNAs among specific lncRNAs, differential lncRNAs, and genes annotated in differential accessible chromatin from HSC differentiated into MEP during erythroid differentiation.

**Supplemental Figure 4.** The expression of lncRNAs with potential regulatory functions at HSC (A, B, C, D, E), MPP (F, G) and CMP (H, I) stages.

**Supplemental figure 5.** (A) Detection of DANCR overexpression in K562 cells by real-time PCR. D1, D2, and D3 represent the cells induced towards erythroid cells with hemin for 1, 2, 3 days, respectively. (B) Expression of globin genes (*HBB, HBE, HBG*) and erythroid genes (*AI.AS2*) after hemin induction with over-expressed DANCR in K562 cells. (C) Overexpression of DANCR promotes hemin induced hemoglobin accumulation in K562 cells. (D-E) Flow cytometry detected the expression of surface maker CD235a^+^-CD71^+^ in K562 cells with overexpression of DANCR and vector control (D) and statistical graphs (E) after hemin induced for 0 day and 3 days. Statistical results were analyzed by student’s t-test and Kruskal-Wallis test, *p < 0.05, **p < 0.01, ***p < 0.001. D0: Day 0, D1: Day 1, D2: Day 2, D3: Day 3, Ctrl: control group, OE: overexpression.

**Supplemental Figure 6. (A)** The chromatin accessibility and the histone modification profile around *ALAS2*.

**Supplemental Table 1**: Motif enrichment results of specific peaks at each stage during erythroid differentiation.

**Supplemental Table 2**: Motif enrichment results of differential peaks during erythroid differentiation.

**Supplemental Table 3**: TFs related to *DANCR* presented in promoter region of *DANCR*.

## Notes

https://www.ebi.ac.uk/ega/dacs/

### Competing Interest Statement

The authors have declared no competing interest.

