## Supplementary figures and images for "The dynamic interactive network of long non-coding RNAs and chromatin accessibility facilitates erythroid differentiation"

### Supplemental Figure 1

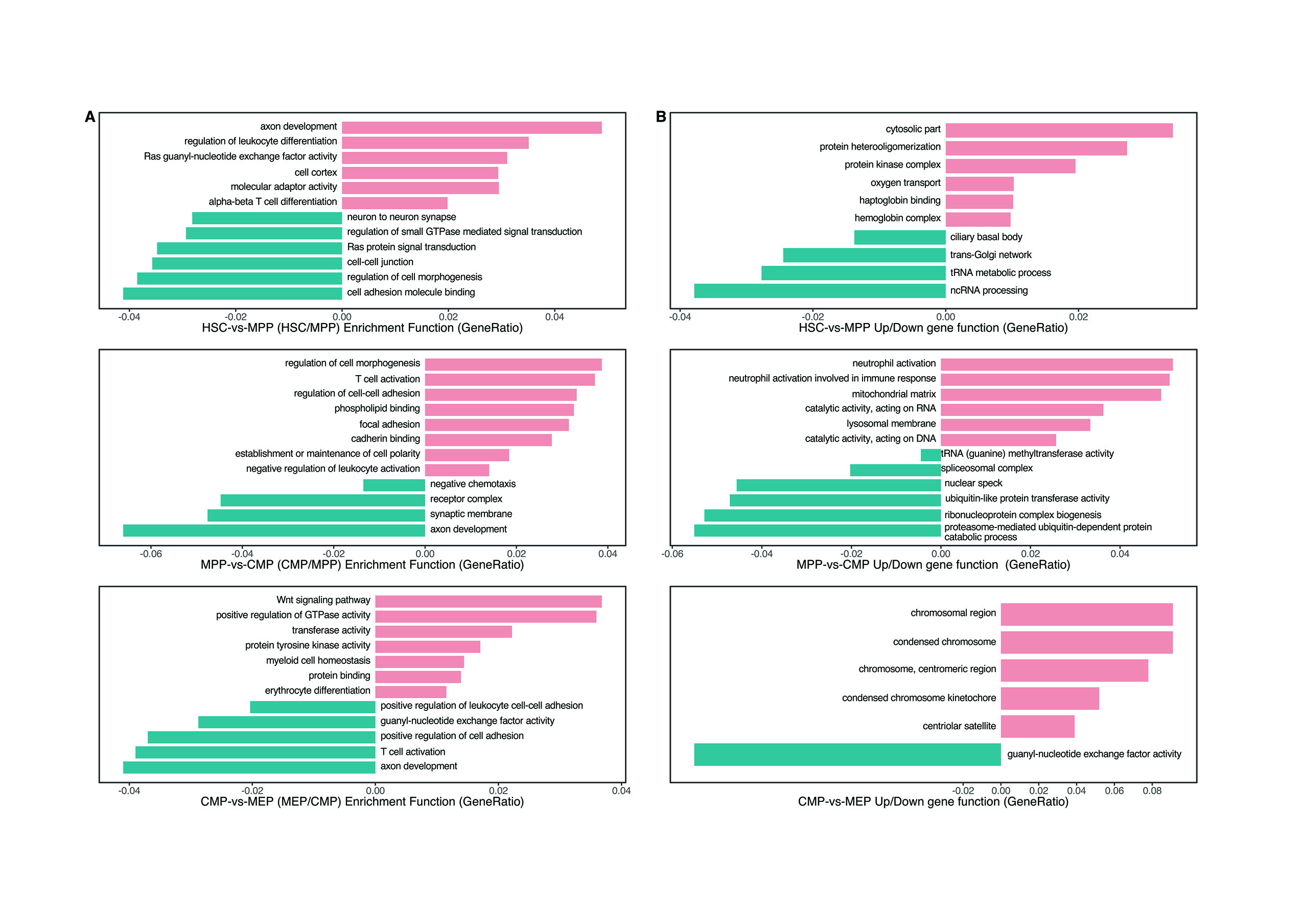

### Supplemental Figure 2

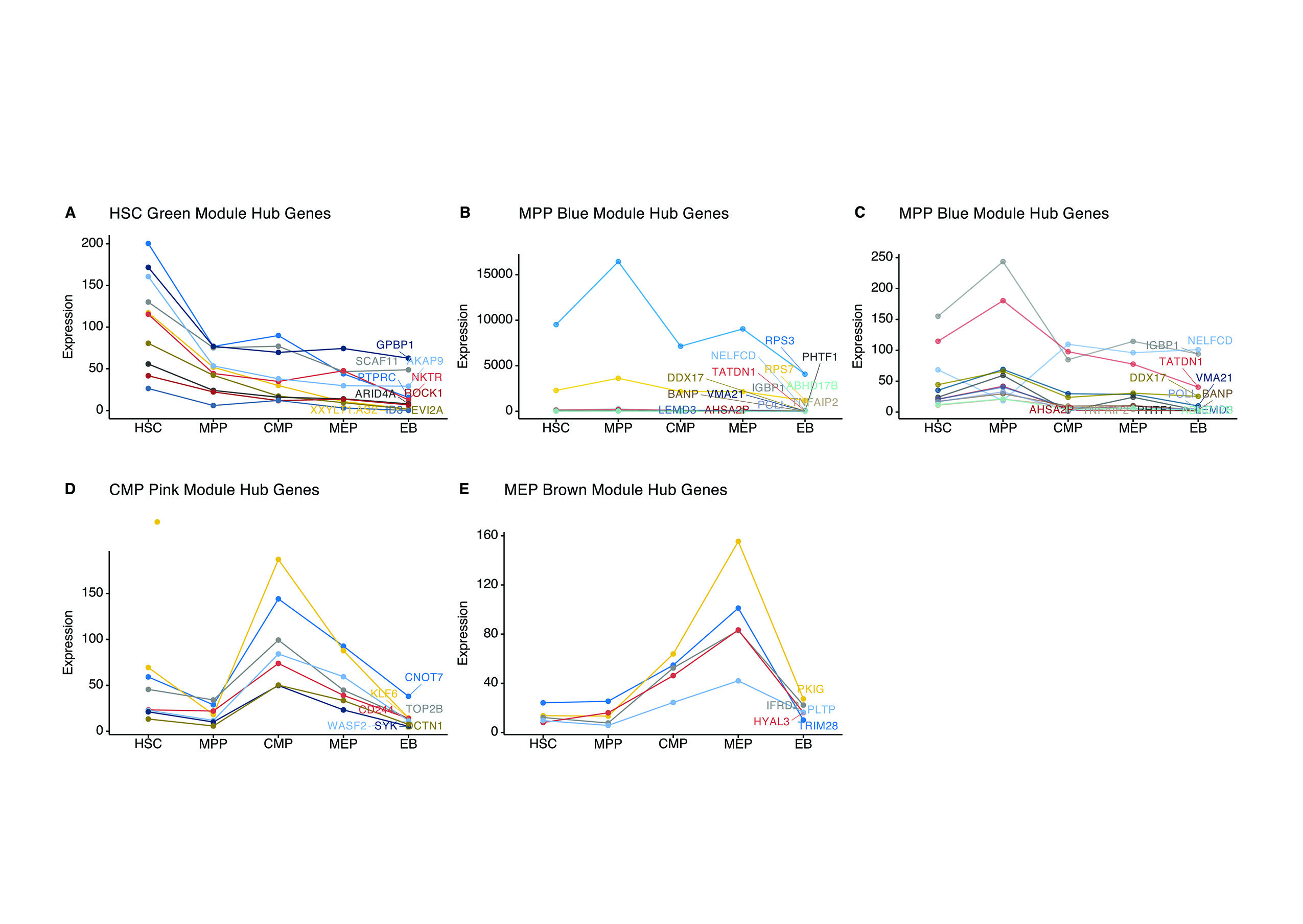

### Supplemental Figure 3

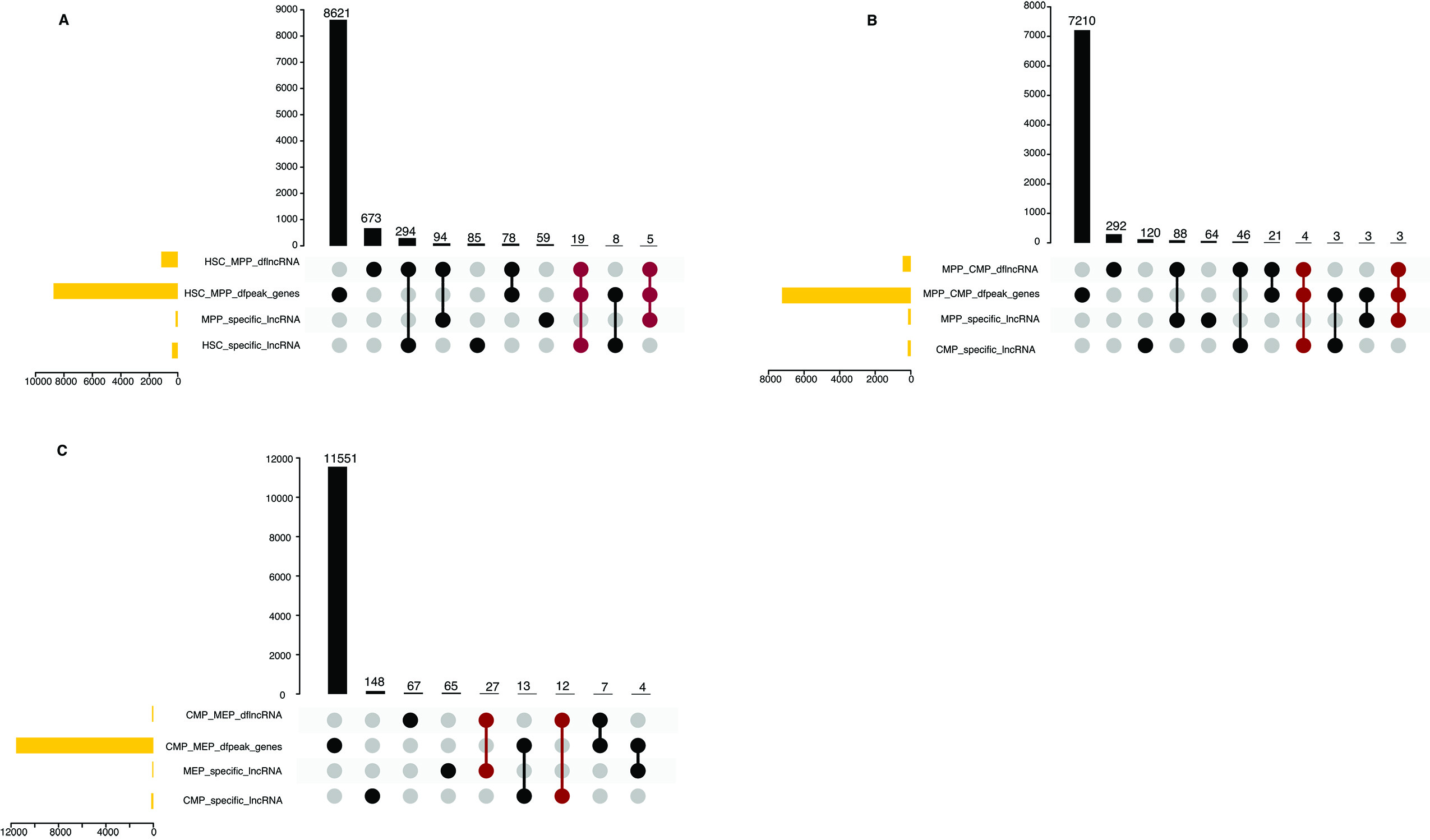

### Supplemental Figure 4

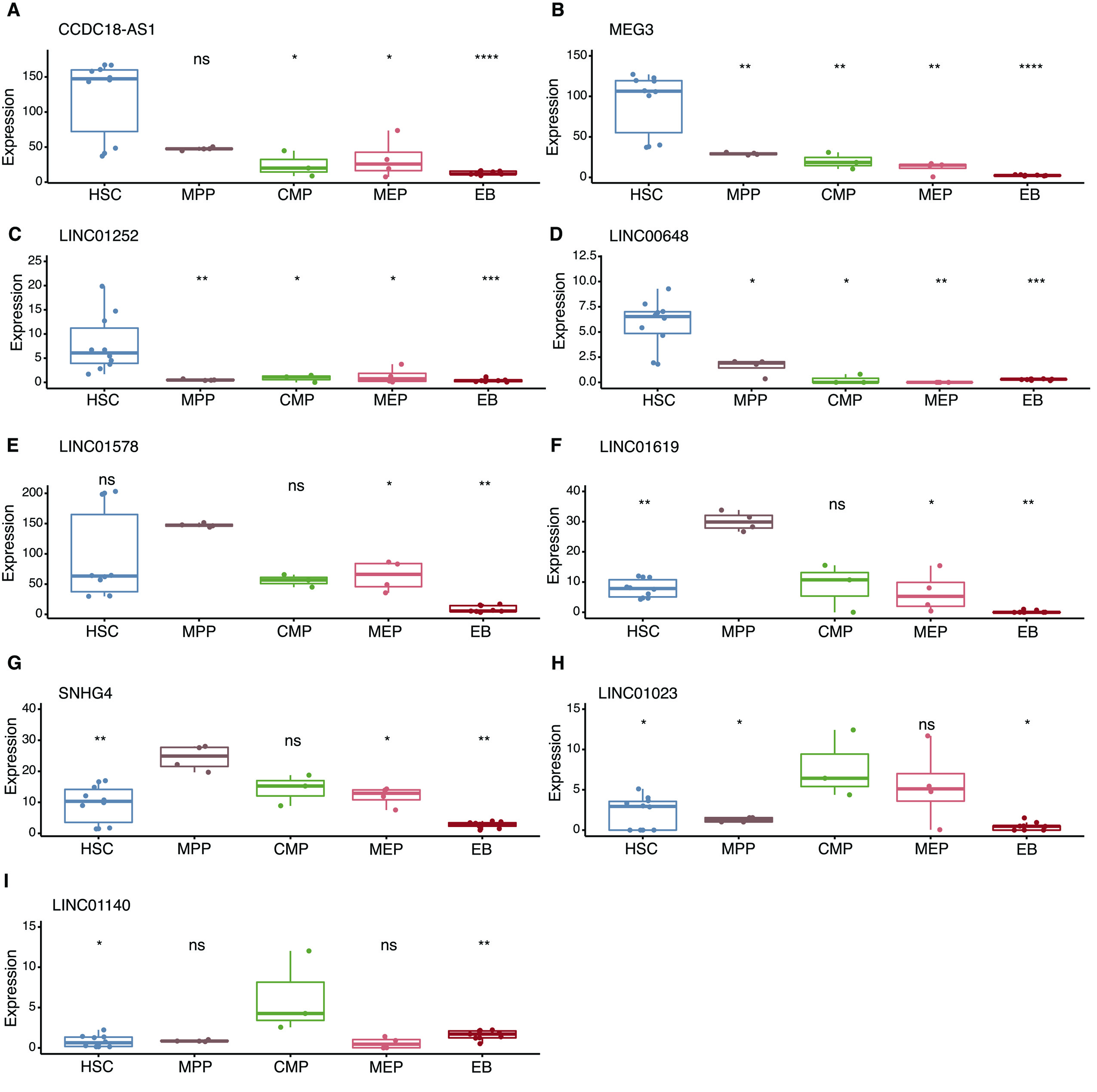

### Supplemental Figure 5

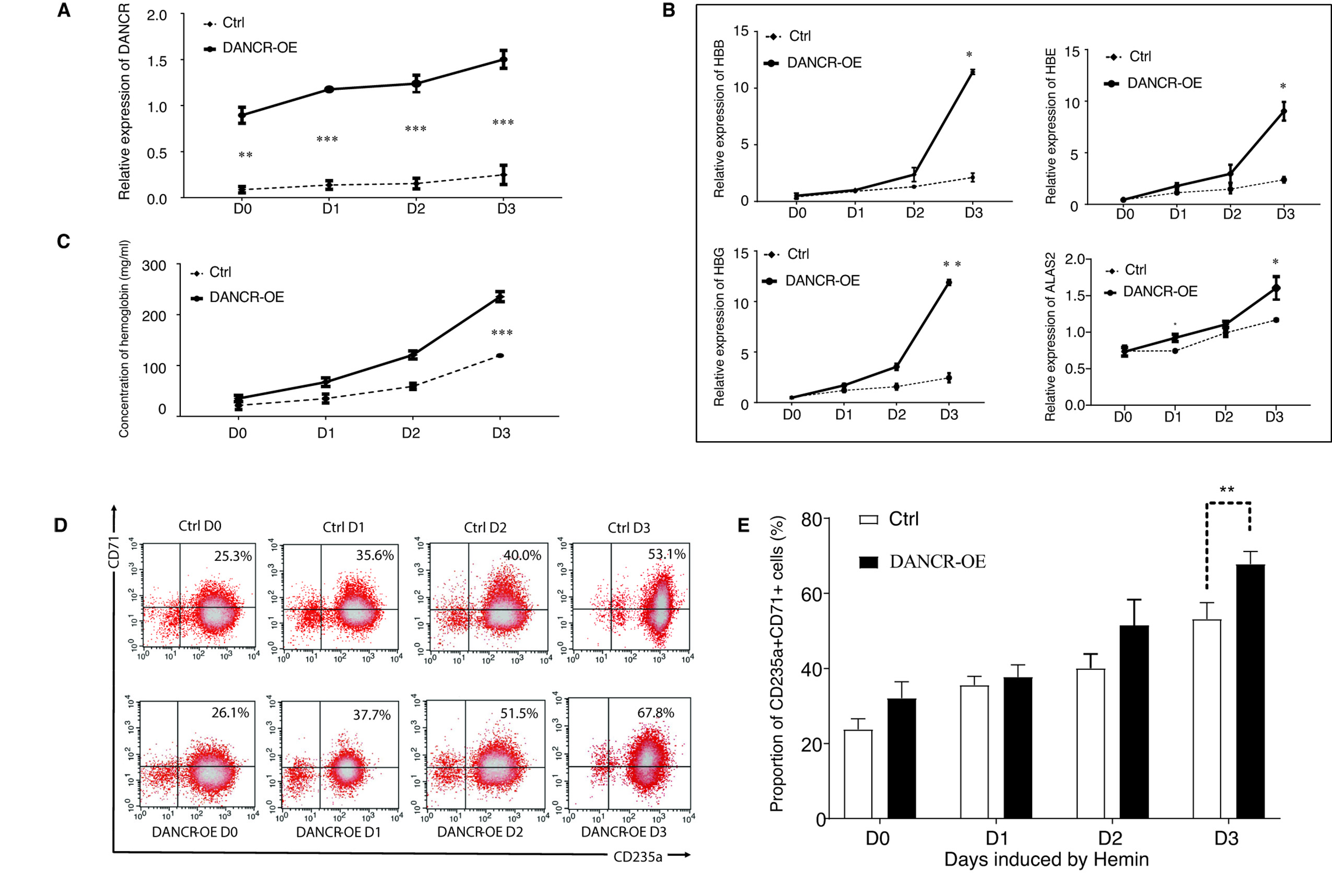

### Supplemental Figure 6

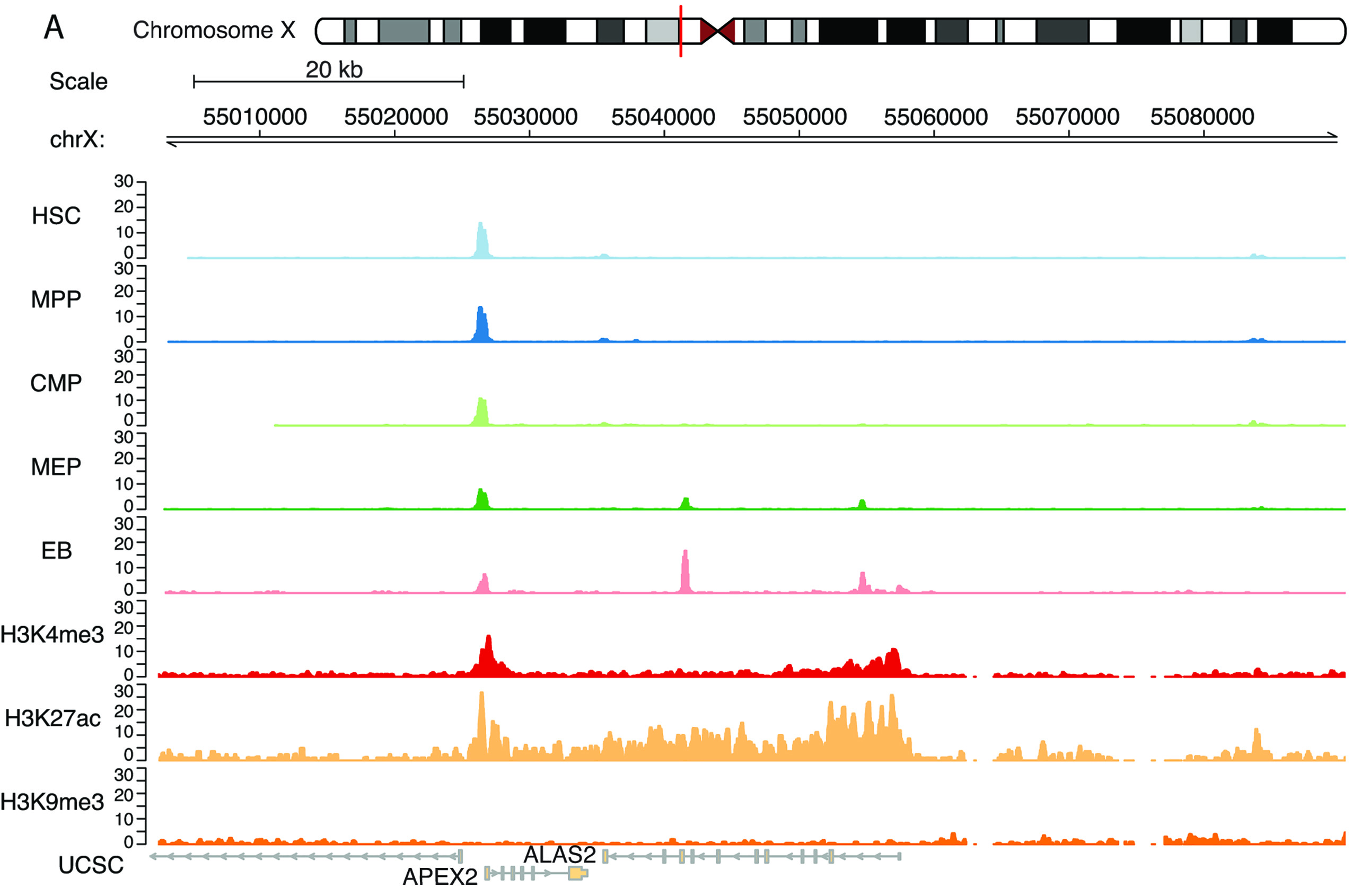
