## Supplemental Table for "The dynamic interactive network of long non-coding RNAs and chromatin accessibility facilitates erythroid differentiation"

**Supplemental Table 1**: Motif enrichment results of specific peaks at each stage during erythroid differentiation.

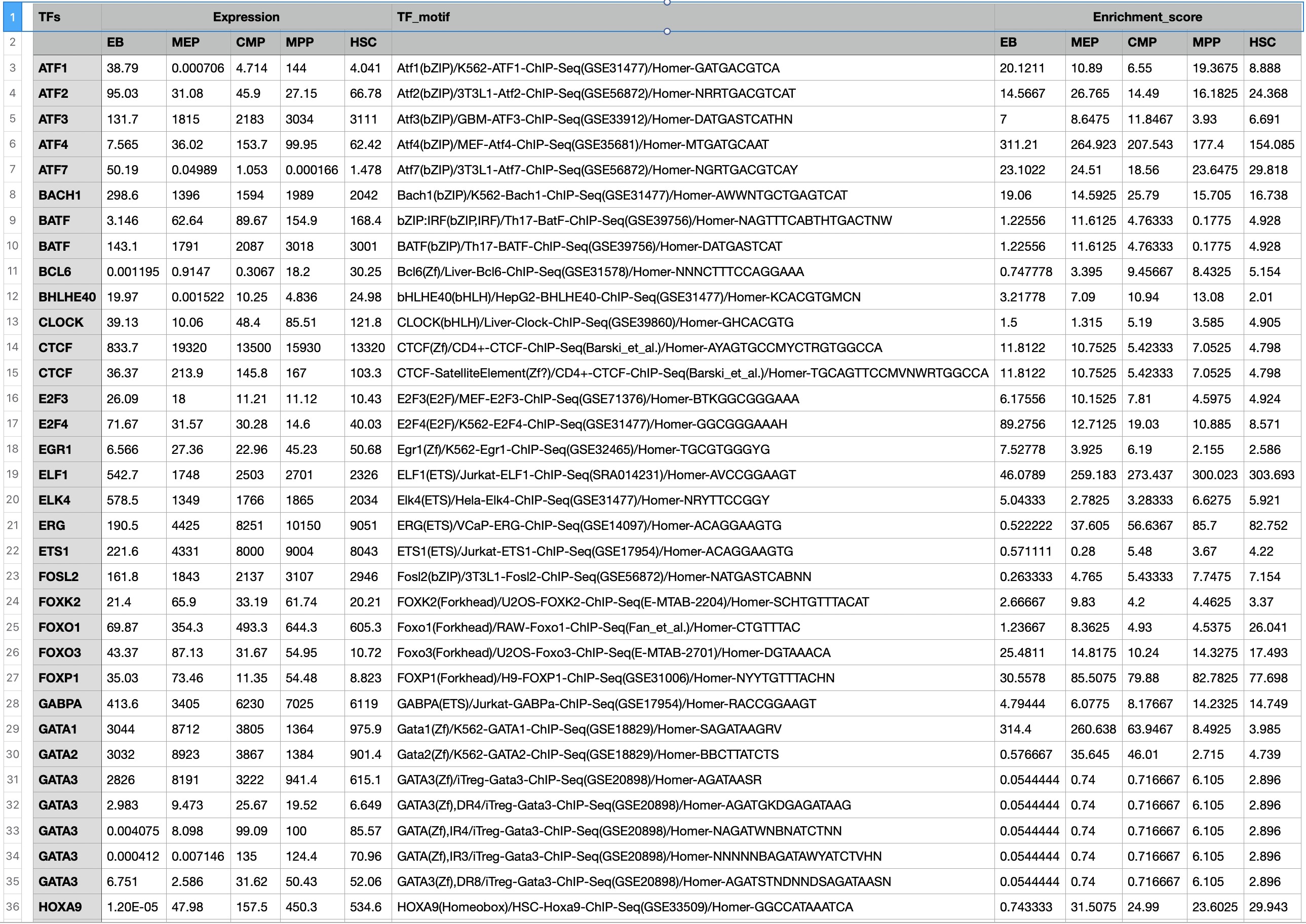

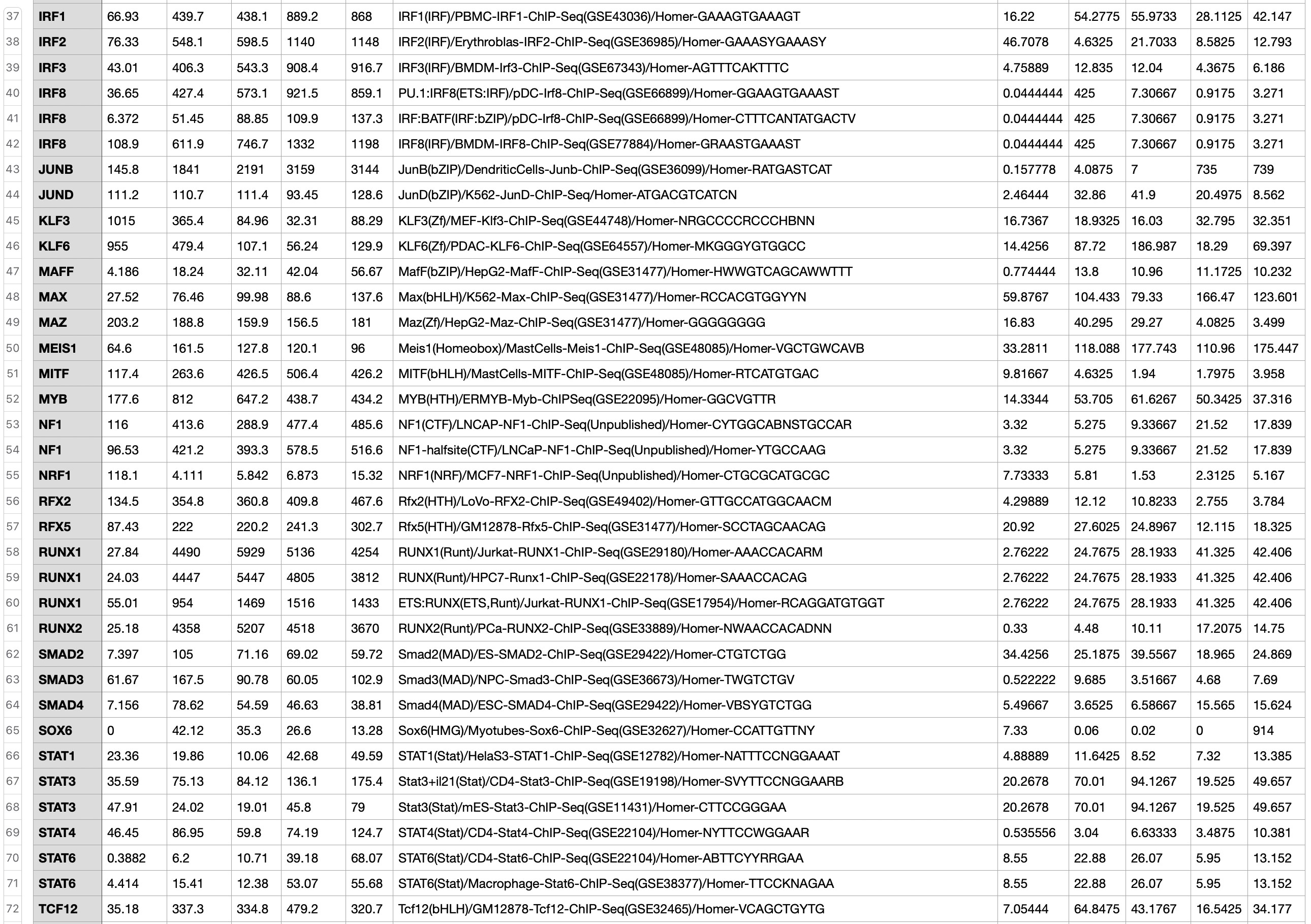

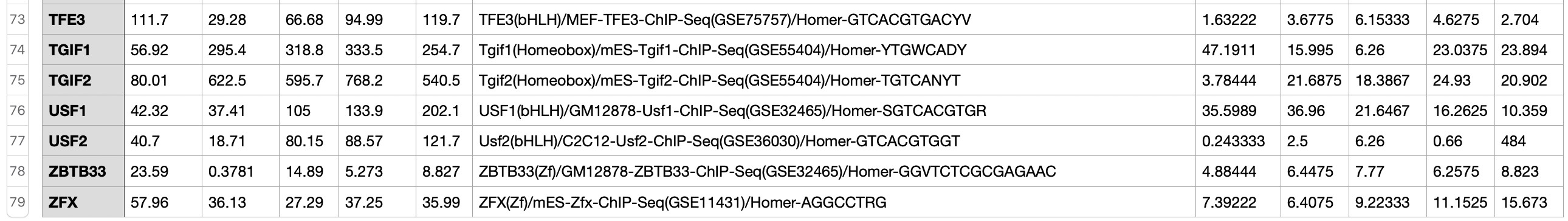

**Supplemental Table 2**: Motif enrichment results of differential peaks during erythroid differentiation.

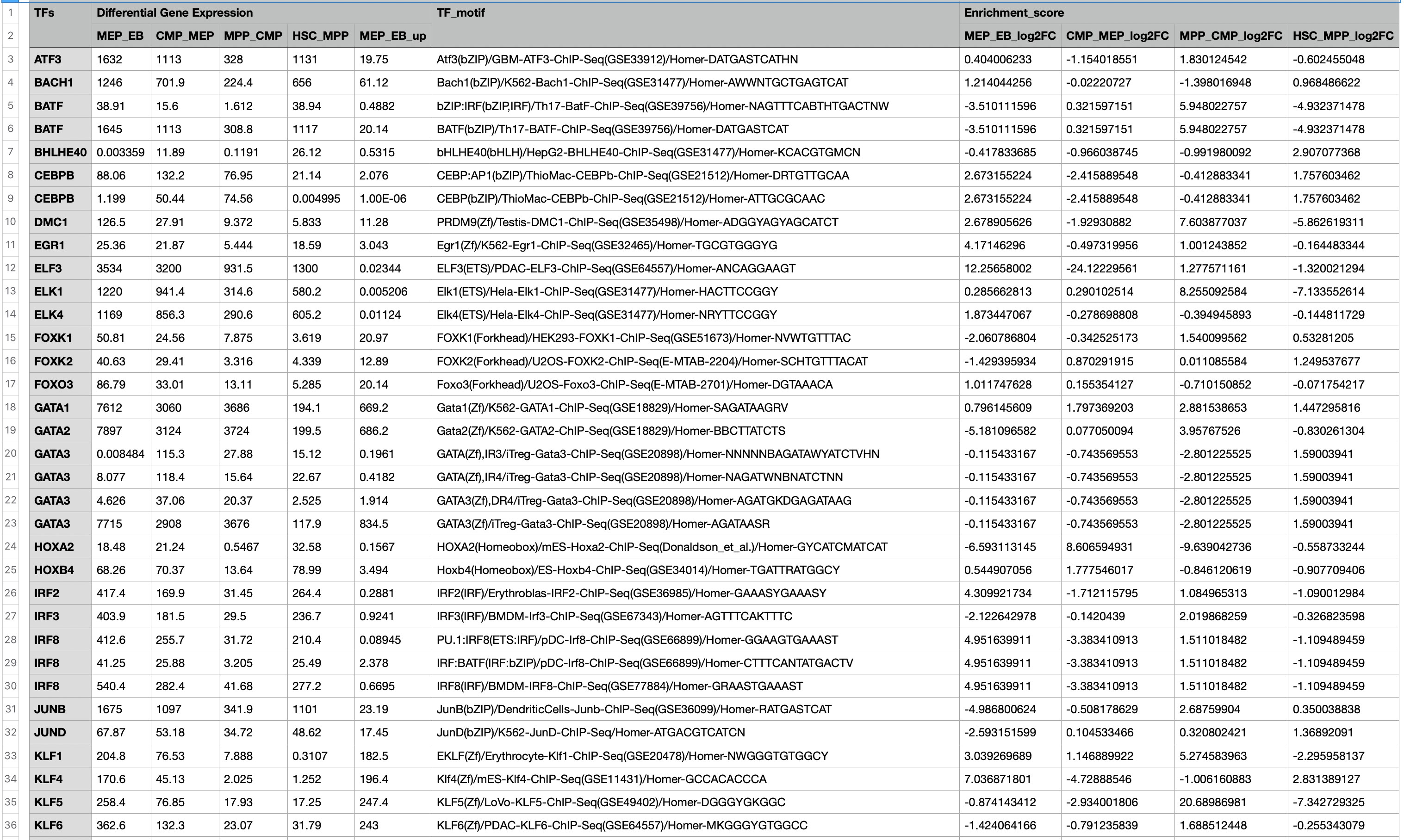

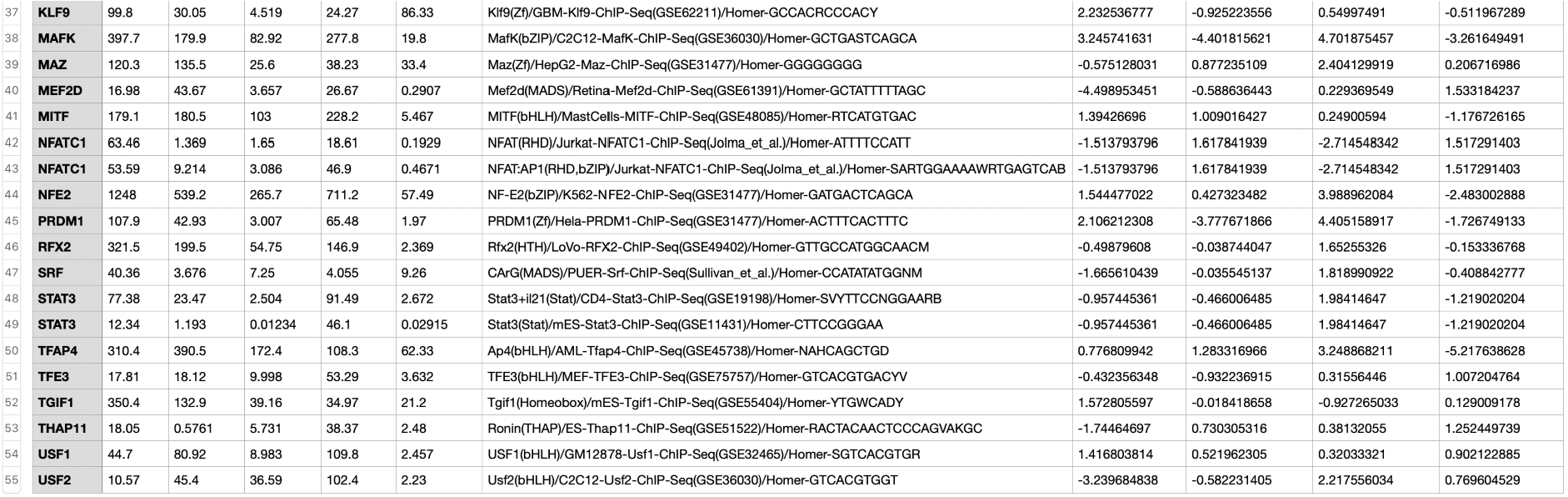

**Supplemental Table 3:** TFs related to *DANCR* presented in promoter region of *DANCR*.

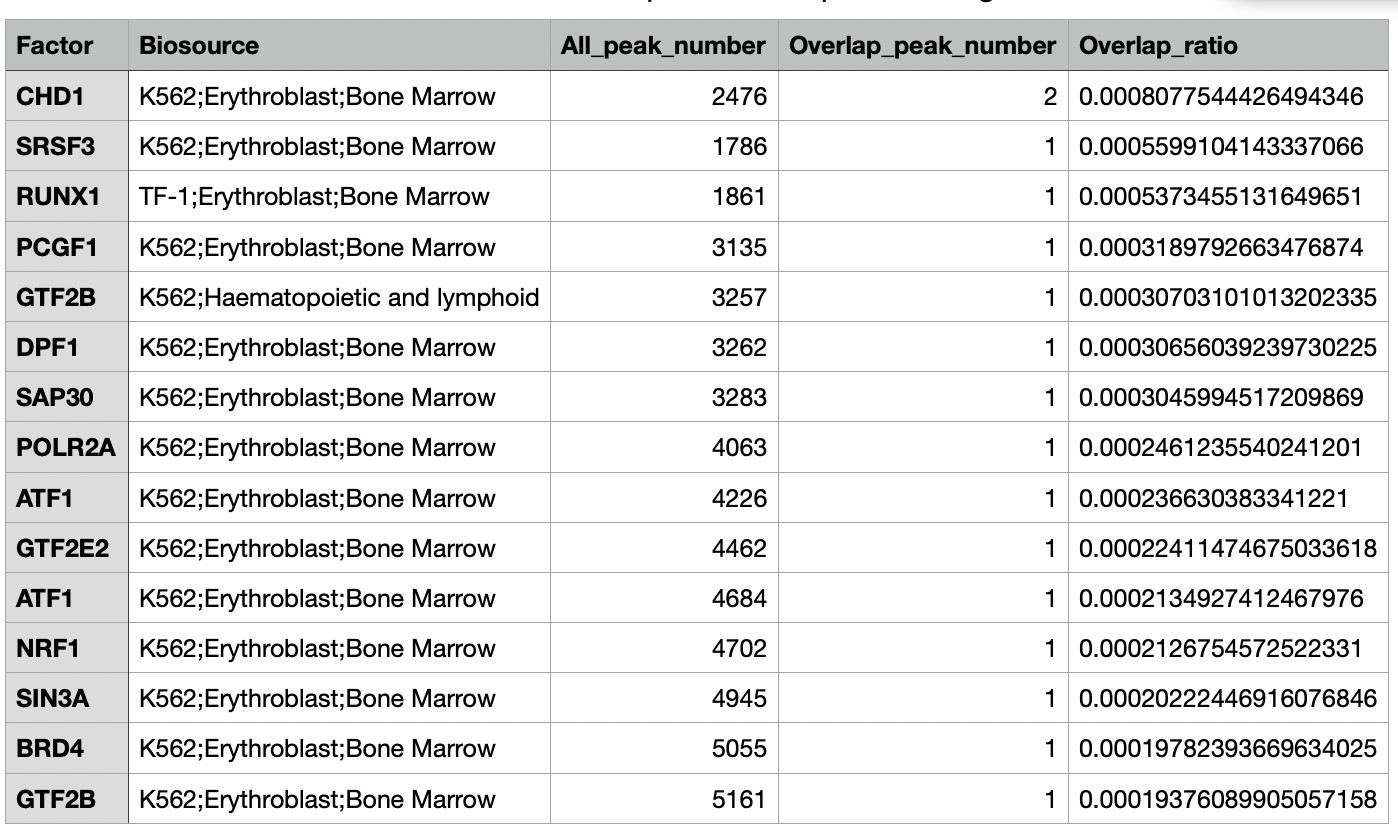
